# A Bispecific Biologic That Robustly Reverses HIV-1 and SIV Latency

**DOI:** 10.64898/2026.10.06.757104

**Authors:** Yaoxing Huang, Manoj S. Nair, Jian Yu, Sue Chong, Hsiang Hong, Hiroshi Mohri, Hiroshi Takata, Benjamin D. Varco-Merth, Alejandra Marenco, Jeffrey D. Lifson, Lishomwa C. Ndhlovu, Anthony Bowen, Lydie Trautmann, Sandhya Vasan, Tae-Wook Chun, Afam A. Okoye, David D. Ho

## Abstract

Antiretroviral therapy suppresses HIV-1 replication but cannot eliminate the latent viral reservoir persisting in memory CD4 T cells. Existing latency reversal candidates have yet to achieve robust viral reactivation at tolerable doses. We engineered T4IL15Δ, a bispecific biologic that selectively delivers IL-15 signaling to memory CD4 T cells, with minimal off-target stimulation. T4IL15Δ reactivated latent virus in 94% (48/51) of clinical samples ex vivo, 75% (15/20) of administrations in HIV-1-infected and ART-suppressed humanized mice, and 100% (6/6) of SIV-infected and ART-suppressed rhesus macaques at doses well tolerated. We also found that off-target stimulation of non-CD4 cells hindered HIV-1 reactivation ex vivo. T4IL15Δ provides a robust reactivation platform that could complement other reservoir-elimination strategies toward an HIV-1 cure.

## Main Text

Despite remarkable success of antiretroviral therapy (ART) in controlling HIV-1 replication in people living with HIV (PLWH), infectious proviruses persist latently within long-lived resting memory CD4 T cells, forming a viral reservoir that remains the major barrier to achieving a cure^1,2^. One conceptually straightforward strategy to purge this reservoir involves two steps: first, reactivating latent viruses, and second, eliminating cells expressing the reactivated viruses. However, achieving robust and frequent latency reversal without systemic toxicity has remained a major challenge.

Numerous candidate latency reversal agents (LRAs) have been investigated but with limited success. Pharmacological approaches, including histone deacetylase (HDAC) inhibitors, disulfiram, and protein kinase C agonists, can perturb HIV-1 latency but have generally produced limited or heterogenous reactivation of the latent reservoir^3–6^. HDAC inhibitors can increase cell-associated HIV-1 transcription in vivo, yet the induction of virus production and release has been limited^4–6^. For example, vorinostat increased HIV-1 RNA in resting CD4 T cells by a mean of 4.8-fold without a significant increase in plasma viremia^4^, while panobinostat increased cell-associated HIV-1 RNA by approximately 3.5-fold but produced only transient and modest increases in plasma viremia^5^. Moreover, these candidate LRAs failed to induce replication-competent viral outgrowth from peripheral blood mononuclear cells (PBMC) at clinically relevant concentrations ex vivo^3^.

Other immune-modulating approaches have also yielded incomplete or inconsistent reactivation. The IL-15 superagonist N-803 alone failed to induce plasma viremia in any HIV-1-infected humanized mice and ART-suppressed SIV-infected macaques^7^. Similarly, the SMAC mimetic AZD5582 induced detectable reactivation in only a subset of SIV-infected macaques after 10 doses^8^. More recently, systemic IL-2 administration showed that robust latency reversal is achievable in most HIV-1-infected, ART-suppressed individuals, but at the cost of substantial systemic toxicity, with severe adverse events requiring hospitalization^9^. Collectively, these studies highlight a central challenge in latency reversal: achieving robust and frequent reactivation of latent proviruses without the toxicity from generalized immune activation.

Once viral reactivation is induced, clearance of infected cells could be pursued through complementary strategies, including broadly neutralizing antibodies^10–13^, induction of CD8 cytotoxic T lymphocytes^14,15^, and/or other reservoir-elimination approaches such as compounds^16,17^ that kill infected cells via the pyroptosis pathway^18,19^.

Here, we report the engineering of a bispecific biologic that selectively delivers a potent activating signal to memory CD4 T cells and robustly and frequently reverses latent HIV-1 ex vivo and in vivo, as well as latent SIV in vivo at doses that were well tolerated. Our studies also show that off-target activation of non-CD4 immune cells impedes HIV-1 latency reversal ex vivo, suggesting that selective immune activation may be a pre-requisite to optimal viral reactivation from CD4 T cells in vivo.

## Results

### Selective targeting of IL-15 to only memory CD4 T cells

Interleukin-15 (IL-15) is a pleiotropic cytokine that supports the survival, proliferation, and functional activation of memory CD4 and CD8 T cells, as well as natural killer (NK) cells^20,21^. IL-15 stimulation rapidly induces expression of the early activation marker CD69 across memory T-cell subsets partly through NF-*κ*B signaling^22^, a pathway shared by HIV-1 activation^23^, suggesting that it could be applicable to latency reversal. Indeed, studies in SIV or SHIV-infected nonhuman primates and people living with HIV-1 have shown that an IL-15 superagonist^24^, N-803, can induce latent reservoir activation under suppressive ART, especially when CD8 T cells are depleted^7,25^. Its clinical utility is constrained, however, by dose-limiting toxicity^26,27^. We therefore pursued a strategy of targeted delivery of IL-15 signaling to only memory CD4 T cells to enhance HIV-1 latency reversal while mitigating off-target activation of memory CD8 T cells and NK cells to minimize toxicity.

We began our effort by constructing a bivalent IL-15-Fc fusion molecule (IL15dsFc) that mimics N-803, but with the use of a disulfide bond to join IL-15 to the “sushi domain” of receptor alpha (R*α*) instead of a linker (**Fig. 1a and Extended Data Fig. 1**). Next, we engineered a bispecific antibody-like molecule termed T4IL15 (**Fig. 1a**), in which one arm of IL15dsFc is replaced by the Fab of ibalizumab, an FDA-approved monoclonal antibody^28^ that targets domain 2 of human CD4 and blocks HIV-1 entry at a step post attachment^29^. As control reagents for our experiments, we also made several constructs, the first of which was T4IL15(Q108D) that contains a point mutation that abolishes IL-15 activity. As a single-arm, non-targeting IL-15 control, we constructed 117/IL15, wherein the Fab of ibalizumab in T4IL15 was replaced by the Fab of an irrelevant monoclonal antibody. As a single-arm ibalizumab control, we utilized 10E8.4/iMab^30,31^, a bispecific virus-neutralizing antibody that lacks IL-15 activity. All five constructs were then tested for CD69 upregulation in peripheral blood mononuclear cells (PBMC) from 12 healthy donors. As expected, T4IL15(Q108D) and 10E8.4/iMab did not show any stimulatory effect compared to unstimulated control (**Fig. 1b**). IL15dsFc and 117/IL15 showed similar activation of memory CD4 T cells, memory CD8 T cells, and NK cells, indicating that valency alone did not contribute to IL-15 potency. In comparison, T4IL15 was modestly more effective in activating memory CD4 T cells, but it also activated memory CD8 T cells and NK cells, which are considered off-target effects for our intended purpose.

**Fig. 1.**
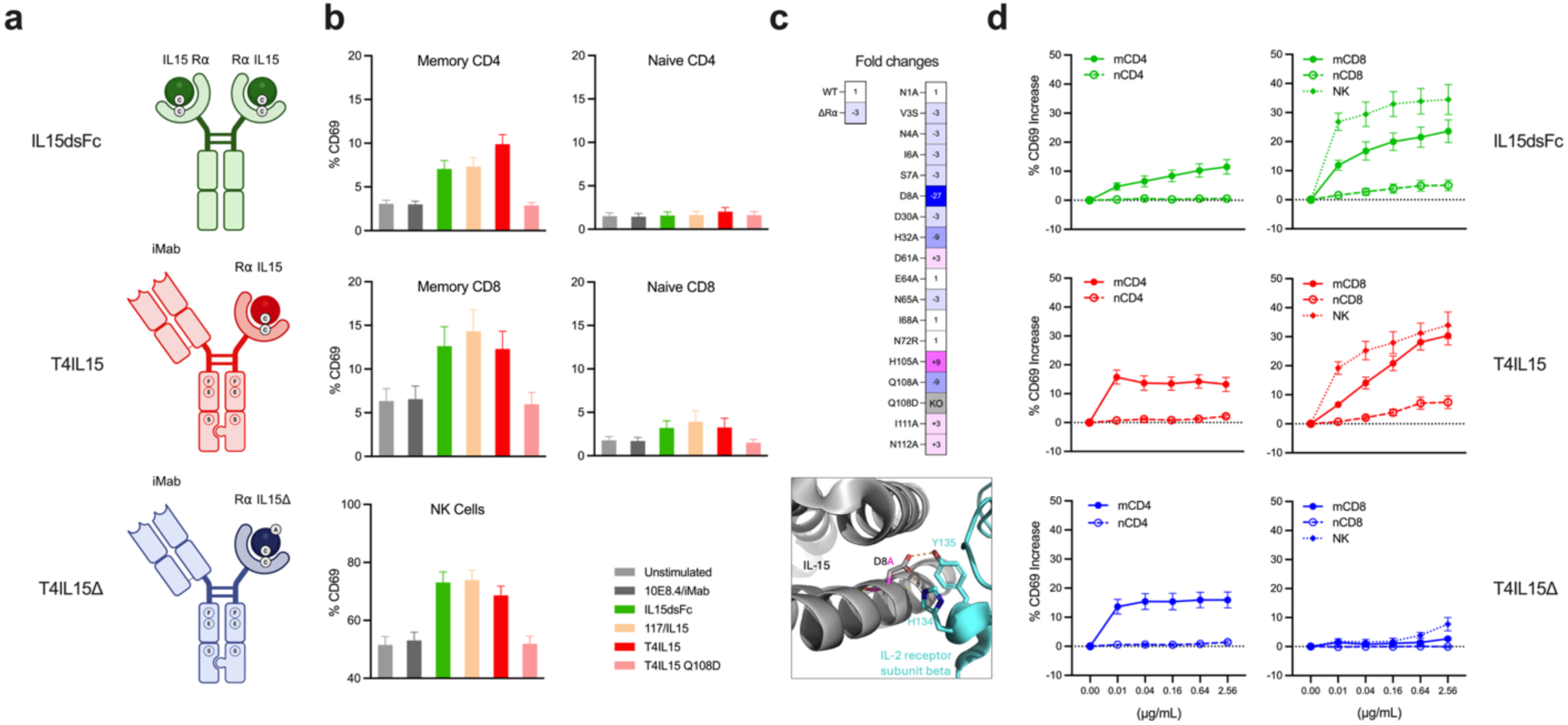
Engineering targeted bispecific biologics. a. Schematic representation of bispecific biologics. The engineered disulfide bond between IL-15 and IL-15Rα is indicated by circled cysteines; the triple mutations to abolish Fcγ receptor interactions are indicated by circled F, E, and S residues in the Fc region; the knob-in-hole heterodimerization design is indicated. b. PBMC activation 24h after treatment with the indicated test agents at 0.1 μg/mL, assessed by CD69 expression. Bars represent mean ± s.e.m. stimulation results (n = 12). c. Heatmap shows relative activity of wild-type and mutant IL-15 proteins to stimulate IL-2Rβ and the common γ-chain. Color scale denotes normalized activity, with blue indicating decreased activity and pink indicating enhanced activity. Structural analysis of the D8A mutation on IL-15 in relation to the receptor is shown, with two predicted hydrogen bond disruptions. d. Dose titration of IL15dsFc, T4IL15, and T4IL15Δ in PBMC stimulation. Cells were treated for 24h across the indicated concentration range, and the increase in activation was quantified by CD69 upregulation in memory CD4, naïve CD4, memory CD8, naïve CD8, and NK cell populations. Points represent mean ± s.e.m. stimulation results (n = 9).

To improve selectivity, we introduced structure-guided mutations at the IL-15 interface with the IL-2/IL-15 β/γ receptor (**Extended Data Fig. 1**). Contact residues predicted to contribute to high-affinity receptor engagement were systematically mutated and screened for IL-15 potency using a reporter assay. Among these variants, the D8A substitution reduced the potency by ∼27-fold relative to wild-type, likely due to the disruption of two key hydrogen bonds (**Fig. 1c**). This mutation was then introduced into T4IL15 to yield T4IL15Δ (**Fig. 1a**), with the expectation that the construct could still engage the β/γ receptors due to an avidity effect of anchoring on the CD4 molecules but could not do so on cells lacking CD4. Indeed, dose-response analysis showed that T4IL15Δ retained robust activation of memory CD4 T cells, while minimizing the stimulation of memory CD8 and NK cells observed for both IL15dsFc and T4IL15 (**Fig. 1d**). Together, these results established T4IL15Δ as a CD4-targeted IL-15 variant that selectively activates reservoir-harboring memory CD4 T cells while avoiding activation of competing immune populations (**Extended Data Table 1**).

### Robust HIV-1 reactivation in PBMC from ART-suppressed individuals

Ex vivo latency reversal assays are conventionally performed using purified CD4 T cells; however, such an approach incompletely captures cytokine-driven regulation of reservoir reactivation within mixed populations of immune cells^32,33^. Specifically, prior studies have shown that IL-15-stimulated CD8 T cells suppressed HIV-1 gene expression across multiple memory CD4 T-cell subsets and that depletion of CD8 T cells enhanced latency reversal in IL-15-based reactivation assays^7,34^. We therefore used PBMC samples to assess whether selective delivery of IL-15 signaling to memory CD4 T cells improved HIV-1 latency reversal ex vivo. Stored PBMC samples from two clinical cohorts of HIV-1-infected, ART-suppressed participants were obtained and tested. The first consisted samples from individuals treated during acute infection while enrolled in the Thai RV254 cohort study^35,36^, and the second consisted of samples from individuals treated at NIH during chronic infection (**Extended Data Table 2**). The test agents included IL15dsFc, T4IL15, and T4IL15Δ, each of which was quality checked by size-exclusion chromatography (**Extended Data Fig. 2**) and assessed at 0.1 *μ*g/mL. Protein kinase activators (phorbol myristyl acetate plus ionomycin or SUW013^37,38^) served as positive controls, and 10E8.4/iMab served as a negative control.

In the first experiment, treatment of PBMC (4 million cells) from 12 RV254 participants with T4IL15 or T4IL15Δ induced robust reactivation of latent HIV-1 in all samples tested (n=12), as evidenced by the detection of virion-associated viral RNA in the culture supernatant 5 days later, whereas exposure to IL15dsFc showed little or no viral reactivation (**Fig. 2a**). In an additional experiment using PBMC (5 million cells) from another 19 ART-suppressed participants from both the RV254 (n=11) and NIH cohorts (n=8), T4IL15Δ induced viral reactivation in 18 of 19 samples tested (**Fig. 2b**). Likewise, treatment of PBMC (6 million cells) from 10 NIH participants with T4IL15 or T4IL15Δ again led to reactivation of latent proviruses (n=10), as evidenced by the release of virion-associated viral RNA into the culture supernatant at frequencies of 60% or 90%, respectively (**Fig. 2c**). Note that the latter experiment was performed at NIH using a slightly modified protocol and a different viral RNA assay. Together, these ex vivo experiments show that our bispecific construct T4IL15Δ can reverse HIV-1 latency in PBMC samples at an unprecedented frequency (39 of 41).

**Fig. 2.**
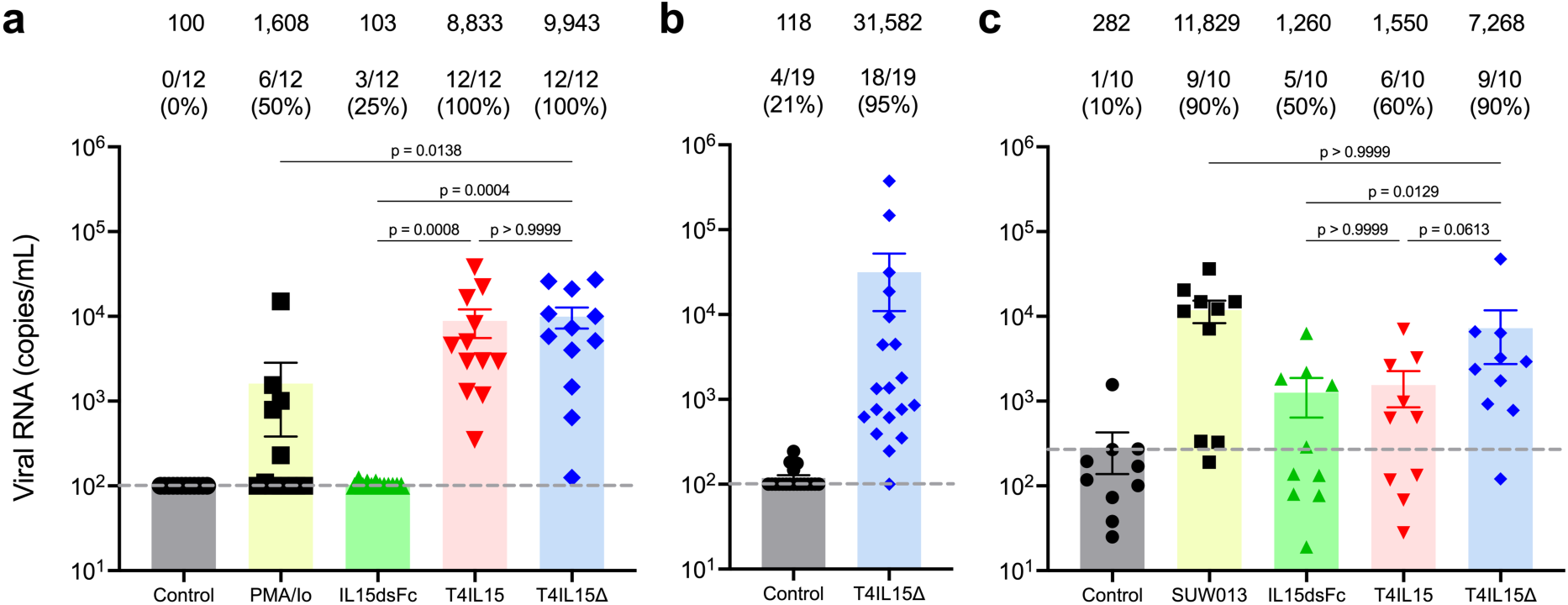
Bispecific biologics induce reactivation of latent HIV-1 from PBMC ex vivo. Reactivation of latent HIV-1 following treatment of (**a**) 4 million PBMC from individuals in the RV254 cohort (n=12), **(b)** 5 million PBMC from individuals in the RV254 and NIH cohorts (n=19) and, (**c**) 6 million PBMC from individuals in the NIH cohort (n=10). Percentage of reactivation is depicted above the bars and statistical significance is shown. Grey dashed line represents LOQ. Mean viral RNA is depicted on the top of each column. Statistical significance was assessed using Dunn’s multiple comparison test in (**a, c**).

### Off target activation of non-CD4 cells impedes HIV-1 latency reversal

Why was IL15dsFc so much less effective in reactivating HIV-1 in PBMC samples ex vivo (**Figs. 2a and 2c)** when it could robustly stimulate memory CD4 T cells (**Fig. 1d**)? To address this apparent discrepancy, we tested IL15dsFc and T4IL15Δ again on PBMC from 10 participants, 5 each from the NIH and RV254 cohorts, as well as on their corresponding CD4-enriched cell populations derived from these samples. The extent of viral reactivation was similar (1.3-fold, but no statistical significance) for IL15dsFc and T4IL15Δ in purified CD4 T cells; however, T4IL15Δ was significantly and markedly better (>10-fold) in whole PBMC samples (**Fig. 3a**). The same set of results plotted by test agents showed a significant and substantial (>7.5-fold) reduction in HIV-1 reactivation by IL15dsFc in PBMC as compared to CD4 T cells (**Fig. 3b**), while reactivation by T4IL15Δ was similar in both cell populations.

**Fig. 3.**
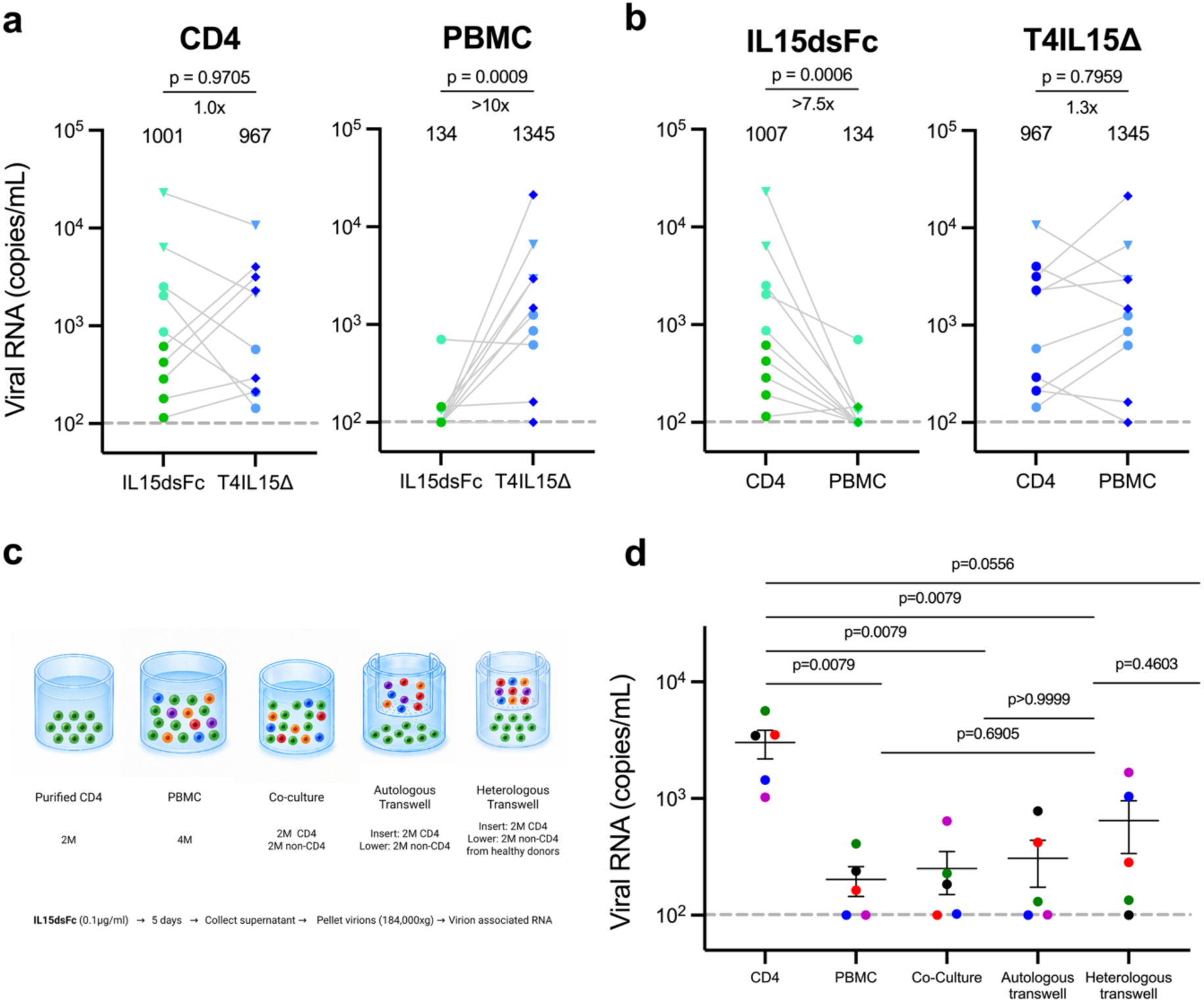
Off-target activation impedes HIV-1 latency reversal. **(a, b)** Comparative reactivation of latent HIV-1 from whole PBMC versus corresponding CD4 T cells enriched by negative selection following treatment with IL15dsFc or T4IL15Δ, each at 0.1 µg/mL (n=10). Viral RNA was purified for quantification from supernatant of 4 million PBMC or 2 million CD4 T cells by ultracentrifugation (circles), or from 6 million PBMC and 3 million CD4 T cells by magnetic particles (inverted triangles), as detailed in Methods. (**c**) Schematic of the five experimental conditions used to assess the effect of non-CD4 cells on HIV-1 latency reversal. (**d**) Comparative reactivation of latent HIV-1 from enriched CD4 T cells, PBMC, CD4 T cells co-cultured with autologous non-CD4 cells, or CD4 T cells in transwell culture with autologous or heterologous non-CD4 cells following treatment with IL15dsFc at 0.1 µg/mL (n=5), as detailed in Methods. Grey dashed line represents LOQ. Statistical significance was assessed using two-tailed Mann–Whitney tests.

These results raised the possibilities that activation of non-CD4 cells by IL15dsFc either impedes HIV-1 latency reversal or leads to the elimination of reactivated reservoir cells. To discriminate between these two scenarios, we conducted experiments using 5 additional RV254 cohort samples under the experimental conditions depicted in **Fig. 3c**. Again, HIV-1 reactivation by IL15dsFc was significantly lower (>14.8-fold) in PBMC compared to enriched CD4 T cells (**Fig. 3d**). As expected, levels of viral reactivation in co-cultures of enriched CD4 T cells with enriched autologous non-CD4 cells were similar to those in PBMC samples. Importantly, when enriched CD4 T cells were co-cultured with enriched autologous non-CD4 cells separated by a transwell membrane, HIV-1 reactivation by IL15dsFc remained similarly low (**Fig. 3d**). This result indicates that cell-to-cell contact is not required for non-CD4 cells to impede latency reversal, making direct killing of reactivated reservoir cells by CD8 T or NK cells unlikely. This conclusion is further supported by the finding that enriched heterologous non-CD4 cells similarly inhibited viral reactivation from participant CD4 T cells by IL15dsFc (**Fig. 3d**), since cells from the healthy donor would not have virus-specific effector cells. Overall, our findings favor the release of soluble factor(s) by activated non-CD4 cells that partially block HIV-1 latency reversal and emphasize the importance of bypassing the activation of these cells to selectively activate the cellular population harboring the latent HIV-1.

### HIV-1 reactivation in infected, ART-suppressed humanized mice

We assessed whether these bispecific constructs could induce reservoir reactivation in vivo. To do so, we used our established humanized mouse model designed to support HIV-1 infection, sustained pharmacologic suppression, and controlled interrogation of latency reversal^13,39^. NSG immunodeficient mice were fully humanized (**Extended Data Fig. 3a**), and then infected with HIV-1 AD8, followed by weekly monitoring until plasma viremia reached a steady state after ∼4 weeks (**Fig. 4a and Extended Data Fig. 3b**), when suppression of HIV-1 replication began with the administration of two bispecific virus-neutralizing antibodies, 10E8.4/iMab297A and VRC07/PGT121 as ART. Once viremia became undetectable, the mice were kept on maintenance therapy with cabotegravir and rilpivirine and attempts to reactivate HIV-1 with each test agent were initiated only after antibody wash-off and three consecutive determinations of aviremia.

**Fig. 4.**
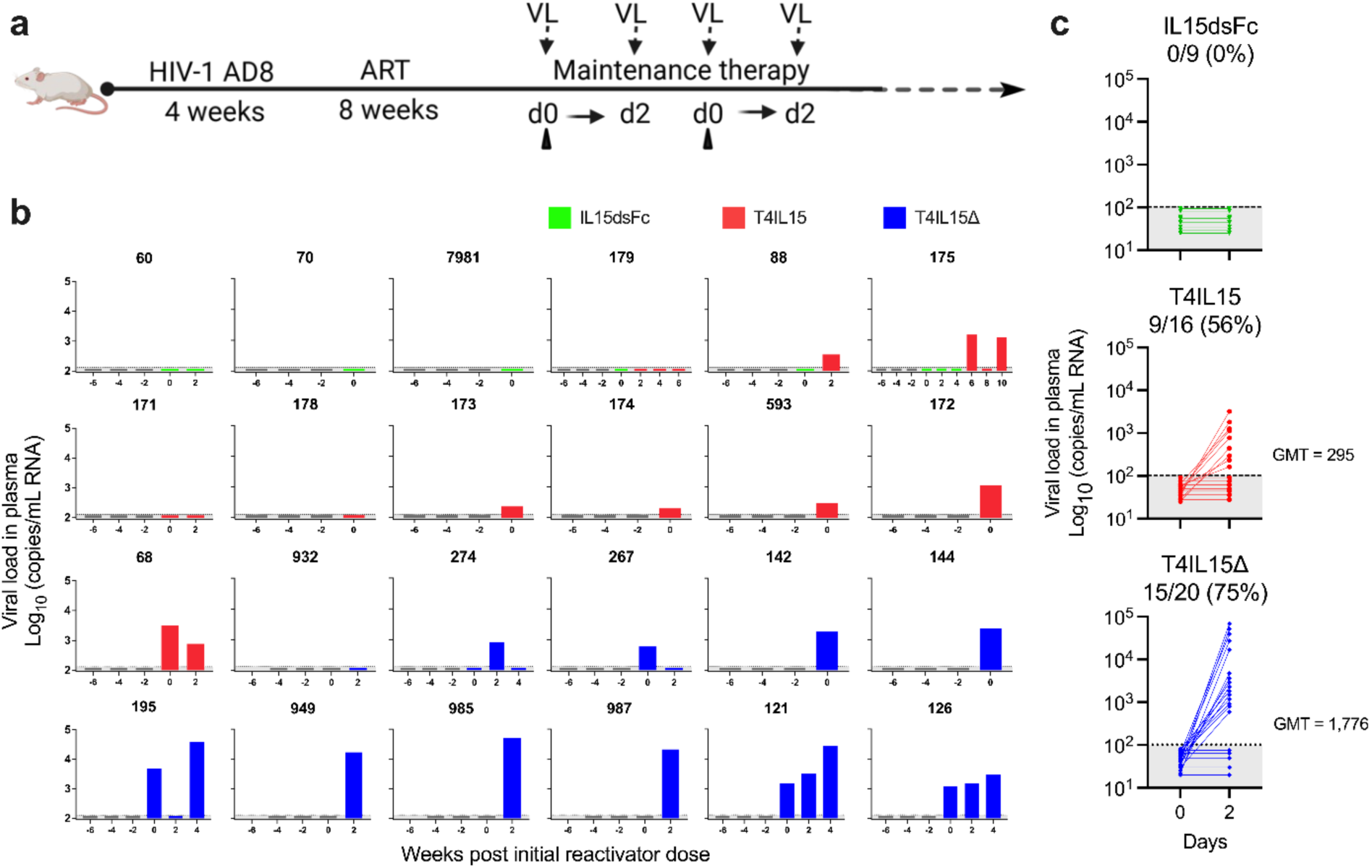
Bispecific biologics induce virion production in ART-suppressed humanized mice. a. Schematic of experimental outline of the in vivo study. Humanized NSG mice were infected with HIV-1 AD8, then treated with two potent bispecific neutralizing antibodies, 10E8.4/iMab297A and VRC07/PGT121, as ART until full suppression. Once viremia was undetectable, mice were kept on maintenance therapy for at least three weeks before initiation of reactivations. Details can be found in the **Methods**. The 297A mutation was made in 10E8.4/iMab to eliminate murine Fc-mediated effector function. b. Viral RNA in plasma samples was measured for each mouse post ART (grey bars). Plasma was collected to measure viral load at day 0 and 48 h after each administration of the test agent. Green bars indicate results for IL15dsFc, red bars for T4IL15, and blue bars for T4IL15Δ. c. Cumulative frequency of in vivo reactivation of HIV-1 detected in plasma of mice on suppressive ART. Dotted lines indicated limit of quantitation. Statistical significance of day 2 viral load differences was assessed using a two-tailed Mann–Whitney test; IL15dsFc to T4IL15 showed p=0.0068, IL15dsFc to T4IL15Δ showed p=0.0009, and T4IL15 to T4IL15Δ showed p=0.0192. Geometric mean titers (GMT) are shown on the right.

While on maintenance therapy, HIV-1-infected and ART-suppressed mice (n = 24) were given a test construct (0.12 mg/kg or ∼3 μg/mouse) intraperitoneally every two weeks, and plasma viremia was measured 48 hours after each administration. IL15dsFc failed to induce any detectable reactivation of the latent reservoir, whereas giving T4IL15 resulted in sporadic detection of plasma viremia (**Fig. 4b**). In contrast, dosing with T4IL15Δ led to frequent detection of plasma viremia, with 3 mice showing evidence of repeated reservoir reactivation following serial administrations. Overall, 75% (15 of 20) of T4IL15Δ doses resulted in detectable plasma viremia, compared to 56% for T4IL15 and 0% for IL15dsFc (**Fig. 4c**). The magnitude of viral reactivation was also significantly higher (6-fold) for T4IL15Δ than for T4IL15. These findings demonstrate that selective CD4-targeted delivery of IL-15 signaling can effectively promote HIV-1 latency reversal in vivo without any observed adverse events.

### T4IL15Δ reactivates SIV in ART-suppressed rhesus macaques

We further investigated T4IL15Δ for SIV latency reversal potential in rhesus macaques. First, we confirmed that the selectivity of our construct was retained in rhesus PBMCs. PBMCs from eight SIV-naïve macaques were cultured for 5 days with IL15dsFc, T4IL15, or T4IL15Δ and assessed for CD69, CD25, and Ki-67 expression. IL15dsFc and T4IL15 induced broad activation of memory CD4 and CD8 T cells, with additional NK-cell stimulation by T4IL15 (**Extended Data Fig. 4**). In contrast, T4IL15Δ preserved dose-dependent induction of CD69, CD25, and Ki-67 in memory CD4 T cells while markedly attenuating responses in memory CD8 T cells and NK cells. Naïve CD4 and CD8 T-cell populations showed comparatively limited responses. Our results confirmed that T4IL15Δ retains similar activity and selectivity in rhesus macaque cells in vitro as observed in human PBMCs, supporting subsequent evaluation of this construct in SIV-infected macaques.

We next evaluated the latency reversal activity of T4IL15Δ in SIV-infected rhesus macaques maintained on suppressive ART. Six monkeys received two intravenous infusions of T4IL15Δ (500 μg/kg) administered 7 days apart, while six contemporaneous animals remained untreated. Both doses of the T4IL15Δ were well tolerated, without any adverse side effects noted. Plasma SIV RNA increased above the assay limit of detection in all six T4IL15Δ-treated macaques following dosing, with recurrent or sustained increases observed after the second infusion, whereas plasma viremia in untreated controls remained largely at or near the limit of detection (**Fig. 5a**). T4IL15Δ induced a clear rise in plasma viremia, with a mean peak level of nearly 10^3^ copies/mL and duration of ∼2 weeks before returning to baseline (**Fig. 5b**). These findings extend our observations from mouse experiments to nonhuman primates, affirming that T4IL15Δ can robustly and frequently achieve latency reversal at doses that are well tolerated in vivo.

**Fig. 5.**
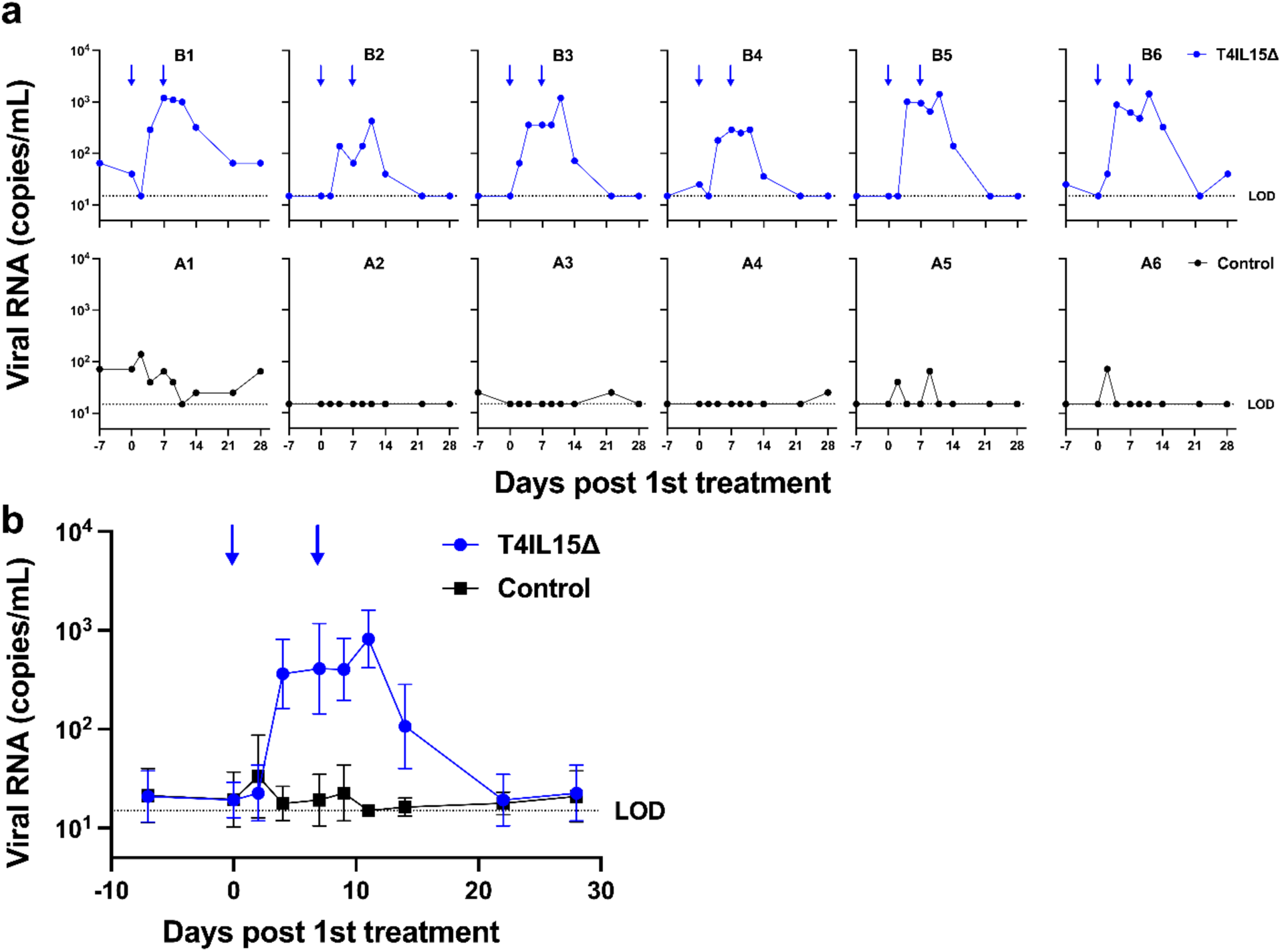
T4IL15Δ induced SIV RNA in plasma of macaques on ART. Plasma viral load profiles of each macaque (**a**) or mean (±SEM) (**b**) in the T4IL15Δ group treated with 2 intravenous infusions at 500 μg/kg (n = 6) versus the non-treated control group (n = 6). Limit of detection was 15 RNA copies/mL of plasma.

## Discussion

Although advances in combination ART have enabled people living with HIV-1 to achieve a near-normal life expectancy, virus eradication remains elusive because replication-competent proviruses persist latently in resting memory CD4 T cells^1,2^. Considerable effort has therefore focused on developing agents to reverse HIV-1 latency, but clinically effective strategies are still limited^40^. A major obstacle is that infectious proviruses reside within an exceedingly small subset of resting memory CD4 T cells that are devoid of known virus-specific markers^40^. Consequently, curative strategies are unable to target only reservoir cells but instead must act on the broader cellular compartment that harbors them. Indiscriminate activation, however, risks immune-mediated toxicity and engagement of other immune cells could counteract latency reversal^7,26,34^.

Here we describe the engineering of a bispecific biologic construct, T4IL15Δ (**Fig. 1a**), that anchors a payload of an attenuated IL-15 in complex with IL-15R*α* to CD4 molecules on the cellular compartment that harbors the latent HIV-1. This novel biologic selectively activated memory CD4 T cells while abrogating off-target activation of CD8 T cells and NK cells (**Fig. 1d and Extended Data Fig. 4; Extended Data Table 1**). In addition, PBMC from ART-suppressed participants from two independent cohorts exposed to T4IL15Δ ex vivo led to latency reversal in 39 of 41 samples, as evidenced by the ex vivo production and release of virion-associated viral RNA (**Fig. 2**). Later, a similar experiment again showed consistent viral reactivation by this construct in 9 of 10 (**Fig. 3a**) PBMC samples. In total, varying degrees of latency reversal were evident in 48 of 51 clinical samples tested ex vivo. Furthermore, HIV-1-infected, ART-suppressed humanized mice given this construct resulted in frequent (75%) detection of plasma viremia 2 days later (**Fig. 4c**). Impressively, T4IL15Δ administrations to SIV-infected macaques resulted in robust levels of plasma viremia in all animals (6 of 6) (**Fig. 5**) while they were maintained on effective antiretroviral therapy, suggesting that the detected virions must be coming from previously integrated proviruses. Taken together, these results demonstrate that our biologic construct could reverse HIV-1 latency ex vivo, as well as HIV-1 or SIV latency in vivo at rates not previously observed^3,7,26,41^. Moreover, we note that latency reversal was achieved at doses of T4IL15Δ that were well tolerated by the animals.

A close examination of the data comparing IL15dsFc to T4IL15Δ (**Figs. 2a and 4c**) leads one to the unavoidable conclusion that off-target stimulation of non-CD4 immune populations must have a negative impact on latency reversal, which was confirmed experimentally (**Fig. 3**) and is also in line with prior reports that activated CD8 T cells could suppress reactivation of latent proviruses^7,34,41^. Experimentally, we showed that activated non-CD4 cells likely impeded latency reversal via soluble factor(s) rather than the killing of reactivated reservoir cells (**Fig. 3d)**. Thus, the effectiveness of T4IL15Δ in latency reversal derives not from a stronger stimulation of CD4 T cells, but from its more selective stimulation. Although T4IL15Δ was initially designed to restrict IL-15 signaling to memory CD4 T cells in order to mitigate toxicity associated with broad immune activation, our findings indicate this selectivity also enhanced the desired biological outcome of activating latent HIV-1 proviruses. This principle may extend beyond IL-15 and should be considered in the development of other latency reversal agents.

To date our study has not addressed the proportion of the HIV-1 latent reservoir that is activated with each administration of T4IL15Δ. Yet, defining that number is essential, because it is obvious that the activated fraction with each administration of any LRA must be sufficiently large so that we are not faced with the prospect of another prolonged intervention. If necessary, T4IL15Δ could be combined with other LRA(s). Moreover, T4IL15Δ could also serve as a vehicle to deliver other agents, such as HDAC inhibitors^4–6^, PKC activators^42^, and SMAC mimetics^8^ to the right target cells to further improve the extent of HIV-1 latency reversal. As a proof of concept, conjugation of T4IL15Δ to romidepsin^6^ synergized memory CD4 T-cell activation compared to the treatment of either agent alone or their combination, while largely preserving cellular selectivity (**Extended Data Fig. 5**).

Ultimately, one major approach to purge the HIV-1 latent reservoir will require both robust activation of latently infected cells and their subsequent elimination. The present study addresses the first of these challenges and demonstrates unprecedented results with our bispecific biologic T4IL15Δ. We also show evidence that its effectiveness depends on its selectivity in activating only memory CD4 T cells, because off-target stimulation of non-CD4 cells impedes latency reversal. Further studies will be needed to determine the optimal application of our construct to the HIV-1 cure effort and how to combine its use with approaches to kill the latently infected cells once they are activated.

## Online Methods

### Data reporting

Power analysis based on guidelines by Institute for Laboratory Animal Research were used to predetermine sample size to estimate minimum number of animals required to detect significant effect of our biologic constructs. The experiments were not randomized, and the investigators were not blinded during experimentation and outcome assessment.

### Ethics statement

The RV254 study (clinicaltrials.gov NCT00796146) was approved by the Institutional Review Boards (IRBs) of Chulalongkorn University in Thailand and the Walter Reed Army Institute of Research in the United States. Leukapheresis products from the NIH were collected from participants in accordance with protocols approved by the Institutional Review Board of the National Institutes of Health (ClinicalTrials.gov ID NCT00039689). Additional human peripheral blood mononuclear cells (PBMCs) from anonymous blood donors were obtained from leukopaks provided by the New York Blood Center, which does not select donors based on gender or race but ensures that all donors are above 18 years of age. Therefore, these PBMCs did not require approval from the Institutional Review Board. All mouse experiments were carried out in strict accordance with the Policy on Humane Care and Use of Laboratory Animals of the United States Public Health Service. The protocol was approved by the Institutional Animal Care and Use Committee (IACUC) at Columbia University (Animal Welfare Assurance no. D16-00003). Virus inoculations were performed under anesthesia, and all efforts were made to minimize animal suffering. Mice were euthanized with CO_2_, with every effort made to minimize suffering. Nonhuman primate work, including experimental protocols, procedures, husbandry and administered agents, was conducted at the Oregon National Primate Research Center (ONPRC) and approved by the ONPRC IACUC (protocol number IP00001053), which is fully accredited by the American Association for Accreditation of Laboratory Animal Care, with an approved Assurance (A3304-01) for the use and care of animals on file with the NIH Office of Laboratory Animal Welfare. The ONPRC IACUC adheres to national guidelines established in the Animal Welfare Act (7 USC Sections 2131–2159) and the Guide for the Care and Use of Laboratory Animals (8th Edition) as mandated by the US Public Health Service Policy, as well as using the standards of the US NIH Guide for the Care and Use of Laboratory Animals (National Academies Press, 2011).

### Construction, expression and purification of biologic molecules

Bispecific antibody-like constructs were generated as previously described^13^. Briefly, four DNA constructs were cloned into the expression vector gWiz: (1) the heavy chain of ibalizumab (MV1HC), containing “hole” mutations and the L234F, L235E, and P331S triple mutations to abolish Fc-mediated effector function; (2) the corresponding ibalizumab light chain (MV1LC); (3) the IL-15Rα sushi domain (73 amino acids; S40C) fused to IgG1 Fc via a triple-GS linker, incorporating “knob” mutations and the L234F, L235E, and P331S mutations in Fc; and (4) IL-15 containing the D8A and L52C mutations.

On day 0, all four constructs were co-transfected into Expi293 cells (ThermoFisher) at 2.5 million cells/mL using polyethyleneimine. Transfected cells were cultured in a shaker incubator at 125 rpm, 37 °C, 8% CO₂, and 80% humidity. On day 5, culture supernatants were harvested, and proteins were purified using Protein A Sepharose (Cytiva) according to the manufacturer’s instruction. T4IL15Δ–romidepsin (T4IL15Δ-RMD) conjugate was generated using a two-step site-specific glycan-conjugation strategy. First, an Fc glycan–modifying proprietary enzyme is used to replace the native Fc-associated carbohydrate on T4IL15Δ with a modified sugar containing a defined azide chemical attachment site. In a second step, romidepsin bearing a complementary cyclooctyne linker is selectively coupled to this site, producing a uniform conjugate while minimizing nonspecific modification of T4IL15Δ. Purified T4IL15Δ-RMD (>90% by SEC) achieved a drug-to-antibody ratio (DAR) of approximately 2 when tested using LC-MS and showed <5% free drug on RP-HPLC. All constructs were validated through size exclusion chromatography for purity **(Extended Data Fig. 2)**, and their activities are summarized in **Extended Data Table 1**.

### In vitro activation of human PBMC

Two million human PBMC from each healthy donor were stimulated with test agents at the indicated concentrations for 18–24 hours. For experiments evaluating T4IL15Δ-RMD, PBMC were treated with T4IL15Δ or T4IL15Δ-RMD at 0.1 µg/mL (equivalent RMD of 3 nM), free RMD at 3 nM, or unconjugated T4IL15Δ (0.1 µg/mL) plus RMD (3 nM) for 24 hours. Following incubation, cells were harvested, washed twice with FACS buffer (1% FBS in PBS), and stained at room temperature for 30 minutes with a cocktail of fluorophore-conjugated antibodies, including CD3–APC (BioLegend), CD4–FITC (BioLegend), CD8–BV421 (BioLegend), CD45RO–APC-Cy7 (BioLegend), CD56–BUV737 (BD Biosciences), CD45RA-PE-Dazzle594 (BioLegend), CCR7-BV650 (BioLegend), and CD69–BV785 (BD Biosciences). Cells were then stained with LIVE/DEAD Fixable Yellow viability dye (Invitrogen) for 10 minutes at room temperature, followed by two additional washes prior to data acquisition on a NovoCyte Penteon flow cytometer (Agilent).

Memory CD4 T cells were defined as CD3^+^CD4^+^CD8^−^CD45RO^+^; naïve CD4 T cells as CD3^+^CD4^+^CD8^−^CD45RO^−^; memory CD8 T cells as CD3^+^CD4^−^CD8^+^CD45RO^+^; naïve CD8 T cells as CD3^+^CD4^−^CD8^+^CD45RO^−^; and NK cells as CD3^−^CD56^+^. In the RMD experiments, CD4 central memory T cells (T_CM_) were defined as CD3^+^CD4^+^CD8^−^CD45RA^−^CCR7^+^, and CD4 effector memory T cells (T_EM_) as CD3^+^CD4^+^CD8^−^CD45RA^−^CCR7^−^. CD69 expression was quantified as the percentage of CD69^+^ cells within each gated population. The percent increase in CD69 expression was calculated by subtracting the percentage of CD69^+^ cells in untreated negative-control samples from that observed after treatment with test agents.

### IL-15 mutagenesis and reporter bioactivity assay

Residues at the IL-15 interface with the IL-2Rβ/γ chains were identified based on the published crystal structure (PDB: 4GS7)^43^. Mutant T4IL15 fusion proteins were generated by co-transfecting expression constructs encoding MV1HC, MV1LC, IL-15Rα–Fc, and individual IL-15 variants into Expi293 cells. Proteins were purified as described above.

Purified T4IL15 mutants were serially diluted across a concentration range of 100 ng/mL to 0.045 ng/mL and evaluated for bioactivity using HEK-Blue IL-2 reporter cells (50,000 cells per well; 96-well plates, 200 µl per well; InvivoGen). Cells were incubated for 24 hours at 37 °C in a humidified CO₂ incubator. Subsequently, 20 µl of culture supernatant was transferred to 180 µl of QUANTI-Blue reagent (InvivoGen) and incubated for 2 hours at 37 °C. Absorbance was measured at 620 nm using a microplate spectrophotometer.

### Reactivation assay for induction of HIV-1 from participant samples ex vivo

Four million or 6 million PBMC from RV254 cohort samples or NIH cohort samples, respectively, were stimulated with each test agent at 0.1 µg/mL for 5 days. 10E8.4/iMab served as a negative control in assays performed at both sites and was used together with IL-2 in samples from the NIH cohort. For positive controls, RV254 cohort samples were treated with PMA (50 ng/mL) plus ionomycin (1 nM) for 24 hours, whereas NIH cohort samples were treated with SUW013 (50 nM) for 48 hours. Since T4IL15 and T4IL15Δ both contain the ibalizumab CD4-binding domain and can block de-novo infection in cultures at concentrations lower than 0.1 µg/mL (FDA label indicates median EC_50_ of 0.008 µg/mL), no additional ART was added to the culture.

Supernatant virion-associated HIV-1 RNA was then isolated prior to quantification. For RV254 samples, virions were first concentrated by ultracentrifugation (180,000 × g, 40 minutes), followed by RNA extraction using the QIAamp DSP Viral RNA Mini Kit (Qiagen) with DNase treatment (Qiagen). RNA copies were quantified by quantitative RT–PCR targeting HIV-1 pol using previously published primer–probe sets and the TaqMan RNA-to-CT One-Step Kit (Applied Biosystems). For NIH cohort samples, virion-associated HIV RNA was quantified using the Cobas 5800 system and HIV-1 Test, version 2.0 (Roche Diagnostics). Comparative statistical analyses were performed using Dunn’s multiple comparison (GraphPad Prism v11.0).

Comparative studies measuring the induction of virion-associated HIV-1 RNA from whole PBMC versus enriched CD4 T cells were conducted using samples from the NIH cohort. CD4 T cells were enriched by negative selection from whole PBMC using EasySep Human CD4+ T-cell isolation kit (StemCell Technologies). Flow through CD4 T cells were assessed for purity using flow cytometry. For comparison, 2-3 million purified CD4 T cells were assessed along with 4-6 million corresponding PBMC samples in parallel. Three sets of samples were analyzed for virion-associated RNA at Columbia, while the remaining two were tested at the NIH, each using methods already mentioned above. Statistical significance was assessed using two tailed Mann-Whitney test.

### Transwell assay to study the suppression of IL15dsFc-induced HIV-1 reactivation in patient samples ex vivo

PBMCs from 5 participants from the RV254 cohort were used for ex vivo co-culture experiments under the conditions depicted in **Figure 3c**. From each PBMC sample, CD4 T cells were enriched by negative selection using EasySep Human CD4+ T-cell isolation kit (Stem Cell Technologies). The corresponding non-CD4 cell fractions were enriched using the negative fraction of the positive selection of human CD4 cells using EasySep Human CD4 T-cell isolation kit. For each autologous co-culture condition, 2 × 10^6^ enriched CD4 T cells were combined with 2 × 10^6^ enriched autologous non-CD4 cells. Enriched CD4 cells (2 × 10^6^) and PBMC (4 × 10^6^) were used as historical controls. For co-culture with non-CD4 cells (2 x 10^6^) from heterologous PBMC of a healthy donor, the cells were enriched negative selection using EasySep Human CD4+ T cell isolation kit (Stem Cell Technologies) before placing in transwell plates with 2 x 10^6^ enriched CD4 T cells from the study participants. Transwell co-cultures were generated by using gamma irradiated tissue culture treated Millicell polystyrene inserts made of translucent PET membrane bearing 0.4 µm pore size. The co-cultures were treated with IL15dsFc at a final concentration of 0.1 µg/mL and maintained for 5 days. Thereafter, culture supernatants were collected and virion associated RNA was extracted following ultracentrifugation of the supernatant at 184,000xg. HIV-1 viral loads in the pelleted virions were quantified by quantitative reverse-transcription PCR (qRT-PCR) in triplicate as described above.

### Humanized mouse studies

All animal experiments were carried out in strict accordance with the Policy on Humane Care and Use of Laboratory Animals of the United States Public Health Service. The protocol was approved by the Institutional Animal Care and Use Committee (IACUC) at Columbia University (Animal Welfare Assurance no. D16-00003; Investigator protocol AABE9556). Humanized NSG mice were generated from CD34^+^ hematopoietic stem cells as previously described by the Columbia Center for Translational Immunology Humanized Mouse Core^13,44^. Briefly, newborn mice (postnatal day 1–5) were irradiated with 100 rad and received intrahepatic injections of 200,000 human CD34⁺ hematopoietic stem cells 6 hours later. Human immune-cell engraftment was assessed 8 weeks after transplantation and monitored longitudinally during viremia sampling using previously described procedures^13,44^. All mice included in subsequent experiments met established thresholds for human immune reconstitution **(Extended Data Fig. 3a)**.

Successfully reconstituted mice were infected with 37,500 TCID_50_ of HIV-1 AD8 (BEI resources, #11346) to establish steady-state infection (**Fig. 4a**). Plasma viral load was measured weekly to assess steady-state levels of infection. Antiretroviral suppression was initiated using a combination of two potent bispecific antibodies, 10E8.4/iMab297A and VRC07/PGT121, dosed weekly at 10 mg/kg. Mice achieving undetectable plasma viremia were subsequently transitioned to maintenance therapy with cabotegravir 400 mg/kg and rilpivirine 600 mg/kg dosed twice per week prior to latency-reactivation studies (**Extended Data Fig. 3b**). During antiretroviral treatment, mice were bled every two weeks to confirm sustained aviremia. Reactivator dosing was initiated only after aviremia had been confirmed.

### In vitro activation of rhesus PBMC

One million PBMC from 8 healthy donor rhesus macaques were stimulated with test agent at the indicated concentrations in RPMI-1640 containing 10% fetal bovine serum and 100 U/ml penicillin-streptomycin (R10) for 5 days. Following incubation, cells were harvested, washed twice with FACS buffer (1% FBS in PBS) and stained at room temperature with a cocktail of fluorophore-conjugated antibodies as previously described^45–47^.

Briefly, PBMCs were first stained with LIVE/DEAD Fixable Near-IR (ThermoFisher Scientific) and biotin-labeled anti-CCR7 mAb (G043H7, biotin; BioLegend) for 30 minutes. The cells were then washed and stained for 30 minutes with combinations of fluorochrome-conjugated surface antibodies, including anti-CD3 (SP34-2, BUV395; BD Biosciences); anti-CD4 (L200, BUV786; BD Biosciences); anti-CD8α (RPA-T8, BV711; BD Biosciences); anti-CD28 (CD28.2, PE-DAZZ; BioLegend); anti-CD95 (DX2, PE; BD Biosciences); anti-CCR5 (3A9, APC; BD Biosciences); anti-CD14 (M5E2, PE Cy7; BioLegend); anti-CD16 (3G8, BV650; BioLegend); anti-CD20 (2H7, Alexa Fluor 700; BioLegend); anti-CD25 (2A3, BUV737; BD Biosciences); anti-CD69 (CH/4, PerCP-Cy5.5; BD Biosciences); and streptavidin BV421. Cells were then washed, lysed/fixed, and permeabilized prior to intracellular staining with anti-Ki-67 (B56, FITC; BD Biosciences) for 45 minutes. Stained cells were analyzed using a FACSymphony A5 (BD Biosciences) and FlowJo software program (version 10.10.0; BD Biosciences).

CD4^+^ and CD8^+^ T cells were divided into memory and naive subsets using criteria previously described^45–47^. Briefly, naïve T cells constitute a uniform cluster of cells with a CD28^moderate^ CD95^low^ phenotype that are CCR7^+^ and CCR5^−^. Memory T cells are a phenotypically diverse memory population that is CD95^high^ or displays one or more of the following non-naive phenotypic features: CD28^−^, CCR7^−^, CCR5^+^. The percent increase in CD69, CD25 and Ki67 expression was calculated by subtracting the percentage of CD69^+^, CD25^+^ or Ki67^+^ cells, respectively, in untreated negative control sample from that observed after treatment with test agent.

### Infection and treatment of rhesus macaques (RMs)

Twelve male rhesus macaques (Macaca mulatta) of Indian genetic background, aged 3-10 years, were used in this study. All RMs were specific pathogen free as defined by being free of Simplexvirus macacinealpha1, simian betaretrovirus, simian T-lymphotropic virus type 1, simian immunodeficiency virus, and *Mycobacterium tuberculosis* at study initiation. All RMs (3-10 years of age) were intravenously inoculated with 1,000 IU of the barcoded SIVmac239M^48^, before starting ART on day 35 post-infection that consisted of daily subcutaneous injections of 5.1 mg/kg/d tenofovir disoproxil, 40 mg/kg/d emtricitabine, and 2.5 mg/kg/d dolutegravir in a solution containing 15% (v/v) kleptose at pH 4.2 as previously described^49^. Following viral suppression, RMs were divided into 2 balanced groups (n=6 each) based on age, major histocompatibility complex 1 (MHC-1) genetics and viral dynamics. After 28 weeks of ART, RMs were treated with either 2 intravenous injections of T4IL15Δ (n = 6) at 500 μg/kg 7 days apart or no treatment (n = 6).

### Quantification of SIV in plasma

SIV RNA levels in plasma were determined using a gag-targeted quantitative real-time/digital RT-PCR format assay, as previously described, providing an assay threshold of 15 SIV RNA copies/mL^50^.

## Data availability

All data supporting the results of this study are available in the article, extended data, and source data files or from the corresponding author upon request.

## Acknowledgements

We thank Zhiteng Li & Yicheng Guo for conducting modeling analysis on Pymol to depict the interface mutations of IL-15 with β/γ receptor. We thank Masahiro Yamashita for kindly providing the Reporter HIV-1 R5 GFP virus. We thank Paul Wender from Stanford University for providing SUW013 to T.W.C. We thank all study participants, Carlo Sacdalan and Somchai Sriplienchan from SEARCH Research Foundation and Chulalongkorn University, and Suteeraporn Pinyakorn from the Henry Jackson Foundation and Military HIV Research Program for providing RV254 samples and related information. We thank the Infectious Disease Resource and Veterinary staff at the Oregon National Primate Research Center for research support and animal care, as well as to the Quantitative Molecular Diagnostics Core of the AIDS and Cancer Program of the Fredrick National Laboratory for the viral load analysis. We also thank L. Picker, O. Ogunwale, W. Goodwin, J. Pacheco-Mendez, S. Hagen, C. Shriver-Munsch, R. MacAllister and R. Bochart for technical advice.

## Funding Statement

This study was supported by funding to D.D.H. from Andrew and Peggy Cherng, Mark and Anla Kingdon, Samuel Yin, Carol Ludwig, and David and Roger Wu, and in part by the Division of Intramural Research of NIAID, NIH to T.W.C. The RV254 study was supported by a cooperative agreement (W81XWH-18-2-0040) between the Henry M. Jackson Foundation for the Advancement of Military Medicine, Inc., and the U.S. Department of Defense. NHP experiments were conducted with funding from the National Institute of Allergy and Infectious Disease (R01MH130197 to LT, AAO and LCN) and by the NIH Office of the Director (P51OD011092) for operation of the Oregon National Primate Research Center. NHP experiments have also been funded in whole or in part with federal funds from the National Cancer Institute, under Contract No. 75N91019D00024 (to JDL).

## Author Contributions

Conceptualization: YH, DDH Resources: AB, LT, SV, TWC, AAO

Investigation and Formal Analysis: YH, MSN, JY, SC, HH, HM, HT, BDV, AM, JDL, TWC, AAO

Funding acquisition: JDL, LCN, LT, TWC, AAO, DDH

Supervision: YH, DDH

Writing – original draft: YH, MSN, HH, DDH with input from all authors

Writing – review & editing: YH, MSN, HH, DDH with input from all authors

## Competing Interests

YH, MSN, JY, HM, and DDH are listed as co-inventors on patent applications filed by Columbia University on the use of described biologics for HIV-1 latency reversal.

## Additional information

Extended data accompanies this manuscript.

