## Supplementary Figure for "A Bispecific Biologic That Robustly Reverses HIV-1 and SIV Latency"

**This file contains**

Supplementary Note
Extended Data Figures 1–5
Extended Data Tables 1–2

**Supplementary Note**

**Disclaimer**

Material has been reviewed by the Walter Reed Army Institute of Research. There is no objection to this publication. The views expressed herein are those of the authors and do not necessarily reflect the official policy or position of the Defense Health Agency, Department of War, the U.S. Government nor the Henry M. Jackson Foundation for the Advancement of Military Medicine. The investigators have adhered to the policies for protection of human subjects as prescribed in AR 70-25. The content of this publication does not necessarily reflect the views or policies of the Department of Health and Human Services, nor does mention of trade names, commercial products, or organizations simply endorsement by the U.S. Government.

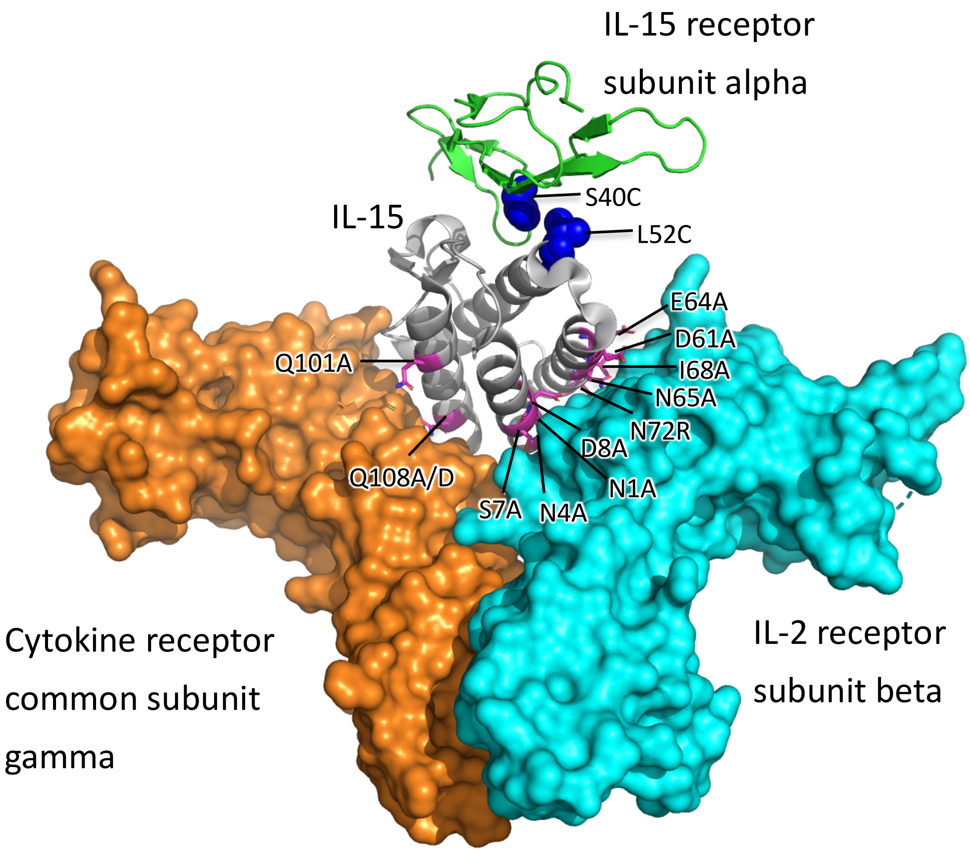


**Extended Data Fig. 1. Structural location of the mutations made and tested in the IL-15–receptor complex (PDB: 4GS7).** Candidate residues evaluated for receptor binding affinity are highlighted in magenta. The disulfide bond linking IL-15 and IL-15 receptor alpha is shown as blue spheres.

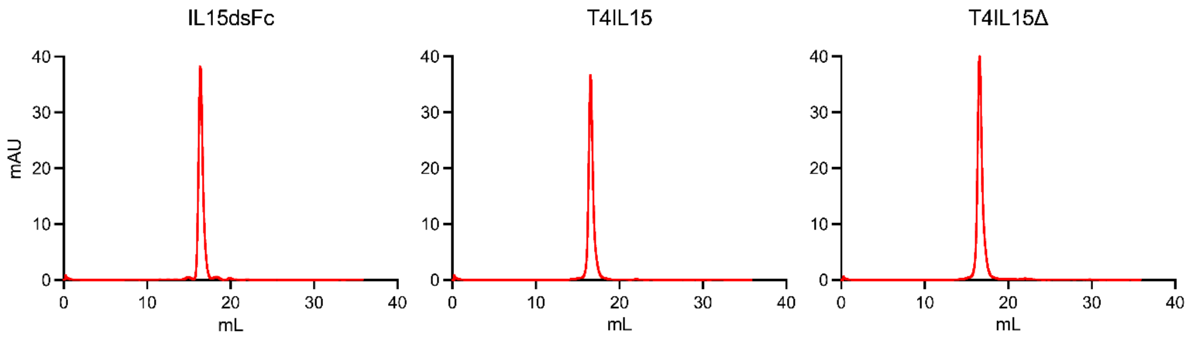


**Extended Data Fig. 2. Size-exclusion chromatography profiles of the engineered bispecific biologics**. All three constructs were validated through size-exclusion chromatography for purity prior to in vitro, ex vivo, or in vivo characterizations.

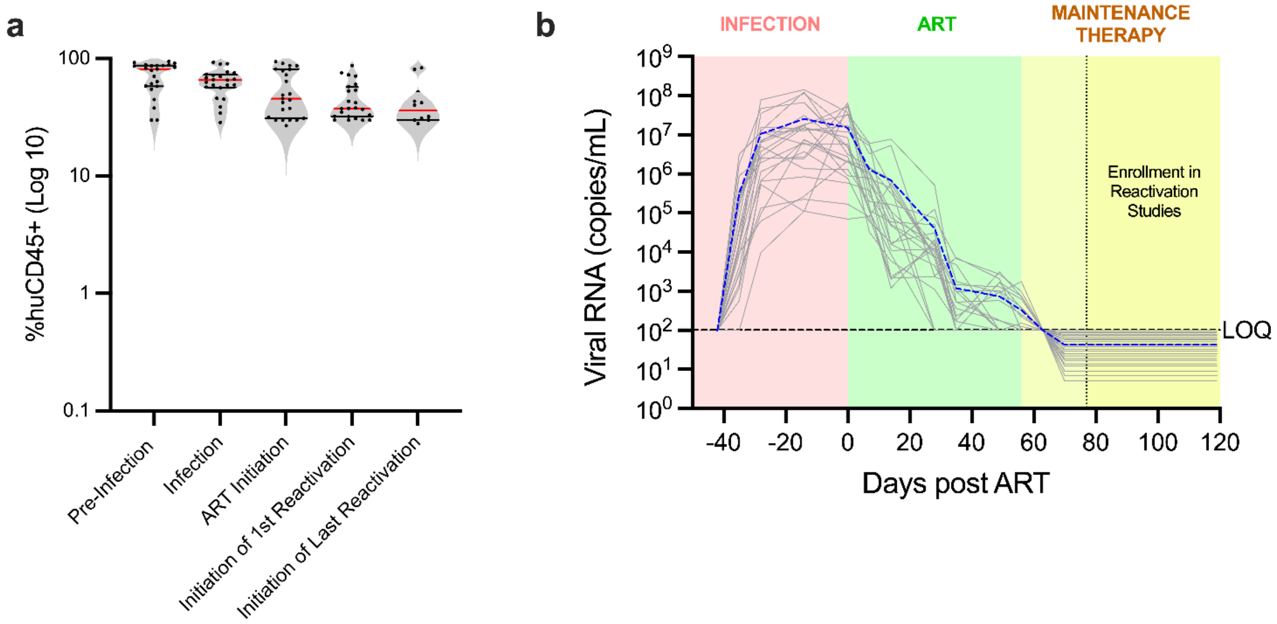


**Extended Data Fig. 3. Humanized NSG mouse model.**

A. Human immune reconstitution levels in humanized mice used in this study. Humanization levels were assessed longitudinally prior to HIV-1 AD8 infection, during the infection period, before ART initiation, during the first reactivation phase, and at initiation of the final reactivation phase. All mice maintained human immune reconstitution above the predefined study thresholds (above 25% of human CD45+ cell engraftment) throughout the experimental timeline.

B. Longitudinal plasma viral loads in humanized NSG mice used in **Fig. 4** (n = 24) across the study timeline, including initial infection (pink), antiretroviral therapy (ART)–mediated suppression (green), and maintenance therapy after suppression (yellow). Black dotted lines indicate the time points at which reactivation studies were initiated following maintenance therapy, defined by at least three consecutive aviremic viral load measurements. Blue dashed line represents the geometric mean. Limit of quantification (LOQ) = 100.

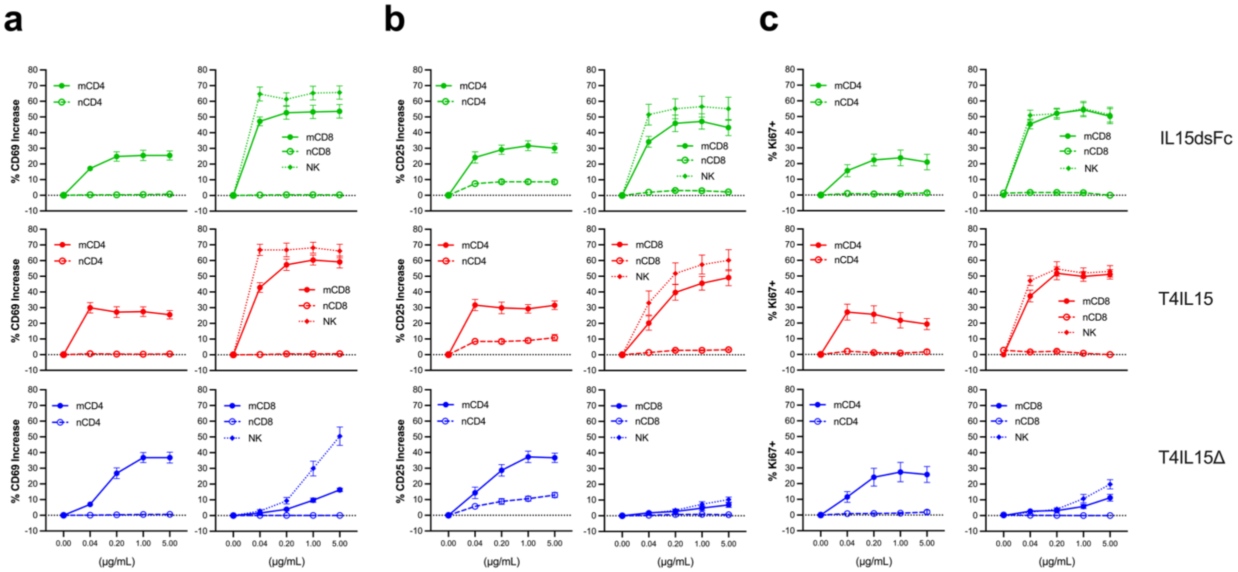


**Extended Data Fig. 4. Engineering targeted bispecific biologics with rhesus PBMCs.** Dose titration of IL15dsFc, T4IL15, and T4IL15Δ in PBMC stimulation. Rhesus PBMCs were treated for 5 days across the indicated concentration range, and the increase in activation was quantified by upregulation of CD69 (A), CD25 (B), and Ki-67 (C) in memory CD4, naïve CD4, memory CD8, naïve CD8, and NK cell populations. Memory CD4 T cells were defined as $CD3^+CD4^+CD95^+$; naïve CD4 T cells as $CD3^+CD4^+CD95^-CD28^+CCR5^-CCR7^+$; memory CD8 T cells as $CD3^+CD8^+CD95^+$; naïve CD8 T cells as $CD3^+CD8^+CD95^-CD28^+CCR5^-CCR7^+$; NK cells as $CD3^-CD20^-CD14^-HLA^-DR^-CD8^{dim}$ cells. Points represent mean ± s.e.m. stimulation results from 8 SIV naïve rhesus macaques.

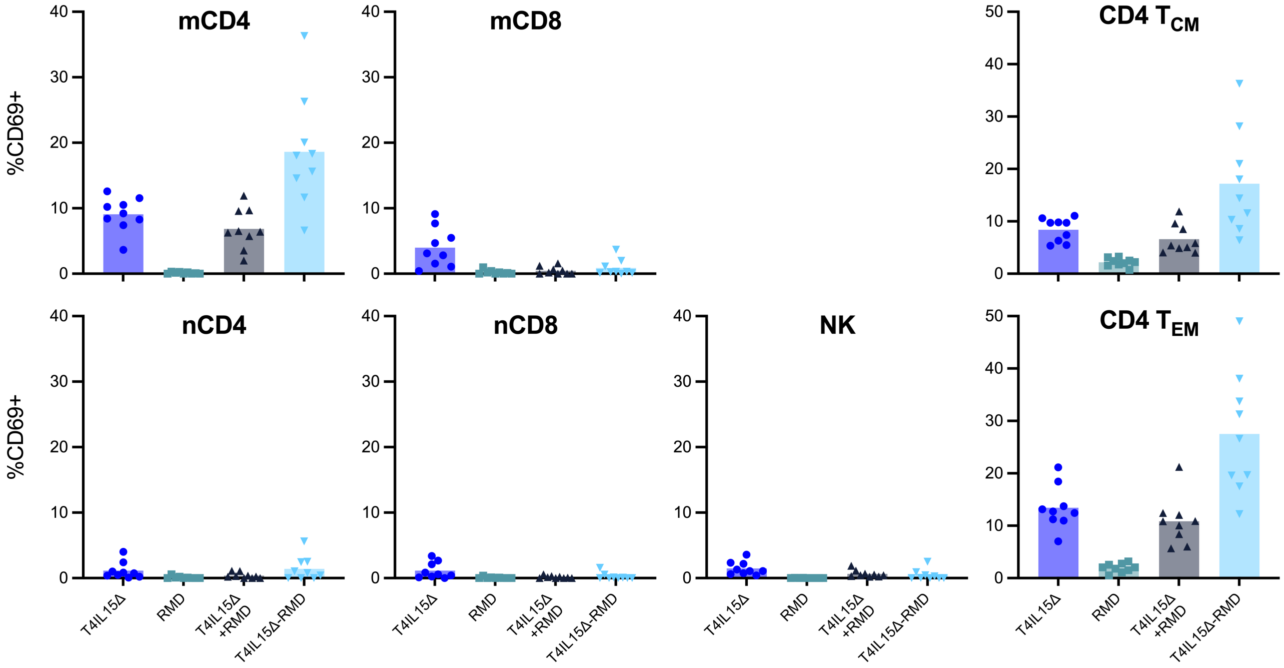


**Extended Data Fig. 5. T4IL15Δ conjugated to romidepsin enhances memory CD4 T-cell activation while largely retaining cellular selectivity.**

Activation of different immune cell populations following treatment of PBMC from healthy donors (n = 9) with T4IL15Δ, romidepsin (RMD), combination of T4IL15Δ plus RMD, or T4IL15Δ chemically conjugated to RMD (T4IL15Δ-RMD). T4IL15Δ and T4IL15Δ-RMD were tested at 0.1 μg/mL, with T4IL15Δ-RMD having a drug-to-antibody ratio (DAR) of 2. RMD was tested at 3 nM, equivalent to the amount of RMD present in 0.1 μg/mL T4IL15Δ-RMD. Cellular activation was assessed by CD69 expression. Each symbol represents an individual donor; bars represent the mean.

| Cell Type | Subset | IL15dsFC | T4IL15 | T4IL15Δ |
|---|---|---|---|---|
| CD4 | Naïve | +/- | +/- | - |
| | Memory | ++ | ++++ | +++ |
| CD8 | Naïve | + | + | - |
| | Memory | + | + | - |
| NK | | + | + | - |

**Extended Data Table 1. Summary of stimulatory activities of biologic constructs.**

| Figure | Cohort | Sample ID | Sex | ART regimen | Time post suppresion (weeks) |
|---|---|---|---|---|---|
| Fig. 2a | RV254 | 2A-1 | M | TDF/FTC/EFV | 76 |
| | | 2A-2 | M | TDF/FTC/EFV | 76 |
| | | 2A-3 | M | TDF/FTC/LPV/RTV | 29 |
| | | 2A-4 | M | TDF/3TC/EFV | 72 |
| | | 2A-5 | M | TDF/3TC/EFV | 72 |
| | | 2A-6 | M | TDF/3TC/EFV | 80 |
| | | 2A-7 | M | TDF/3TC/EFV | 80 |
| | | 2A-8 | M | ABC/3TC/DTG | 71 |
| | | 2A-9 | M | ABC/3TC/DTG | 88 |
| | | 2A-10 | M | ABC/3TC/DTG | 88 |
| | | 2A-11 | M | TDF/3TC/EFV | 88 |
| | | 2A-12 | M | TDF/3TC/EFV | 88 |
| Fig. 2b | RV254 | 2B-1 | M | TDF/FTC/EFV | 16 |
| | | 2B-2 | M | TDF/3TC/LPV/RTV | 94 |
| | | 2B-3 | M | TDF/3TC/RAL | 88 |
| | | 2B-4 | M | TDF/FTC/EFV | 40 |
| | | 2B-5 | M | TDF/3TC/RAL | 88 |
| | | 2B-6 | M | TDF/FTC/EFV | 40 |
| | | 2B-7 | M | TDF/FTC/EFV | 40 |
| | | 2B-8 | M | TDF/FTC/EFV | 11 |
| | | 2B-9 | M | TDF/FTC/EFV | 11 |
| | | 2B-10 | M | TDF/FTC/EFV | 11 |
| | | 2B-11 | M | TDF/3TC/EFV | 41 |
| | NIH | 2B-12 | M | CAB/RPV | 4.7 |
| | | 2B-13 | M | DRV/COBI/FTC/TAF | 6.8 |
| | | 2B-14 | M | BIC/FTC/TAF | 11.7 |
| | | 2B-15 | M | BIC/FTC/TAF | 9.2 |
| | | 2B-16 | M | DRV/ABC/3TC | 17.5 |
| | | 2B-17 | M | DRV/RPV | 15.2 |
| | | 2B-18 | M | BIC/FTC/TAF | 1.6 |
| | | 2B-19 | M | BIC/FTC/TAF | 15.2 |
| Fig. 2c | NIH | 2C-1 | M | CAB/RPV | 4.7 |
| | | 2C-2 | M | DRV/COBI/FTC/TAF | 6.8 |
| | | 2C-3 | M | BIC/FTC/TAF | 11.7 |
| | | 2C-4 | M | BIC/FTC/TAF | 9.2 |
| | | 2C-5 | M | DRV/ABC/3TC | 17.5 |
| | | 2C-6 | M | DRV/RPV | 15.2 |
| | | 2C-7 | M | BIC/FTC/TAF | 1.6 |
| | | 2C-8 | M | BIC/FTC/TAF | 15.2 |
| | | 2C-9 | M | DRV/3TC | 13.6 |
| | | 2C-10 | M | DRV/ABC/3TC | 15.2 |
| Fig. 3a/b | RV254 | 3A-1 | M | TDF/3TC/EFV | 25 |
| | | 3A-2 | M | TDF/3TC/EFV | 24 |
| | | 3A-3 | M | TDF/3TC/EFV | 24 |
| | | 3A-4 | M | TDF/3TC/EFV | 11 |
| | | 3A-5 | M | TDF/3TC/EFV | 24 |
| | NIH | 3A-6 | M | BIC/FTC/TAF | 11.7 |
| | | 3A-7 | M | BIC/FTC/TAF | 9.2 |
| | | 3A-8 | M | DRV/ABC/3TC | 17.5 |
| | | 3A-9 | M | DRV/3TC | 13.6 |
| | | 3A-10 | M | DRV/ABC/3TC | 15.2 |
| Fig. 3d | RV254 | 3D-1 | M | TDF/3TC/DTG | 233 |
| | | 3D-2 | M | ABC/3TC/DTG | 103 |
| | | 3D-3 | F | ABC/3TC/DTG | 93 |
| | | 3D-4 | M | ABC/3TC/DTG | 91 |
| | | 3D-5 | M | TDF/3TC/DTG | 94 |

**Extended Data Table 2. Demographic data for cohorts that provided PBMC samples.** ART abbreviations: TDF, tenofovir disoproxil fumarate; FTC, emtricitabine; EFV, efavirenz; LPV, lopinavir; RTV, ritonavir; 3TC, lamivudine; DTG, dolutegravir; RAL, raltegravir; CAB, cabotegravir; RPV, rilpivirine; DRV, darunavir; COBI, cobicistat; TAF, tenofovir alafenamide; BIC, bictegravir; ABC, abacavir.
